# improving diagnosis and monitoring of treatment response in pulmonary tuberculosis using the molecular bacterial load assay (MBLA)

**DOI:** 10.1101/555995

**Authors:** Wilber Sabiiti, Khalide Azam, Davis Kuchaka, Bariki Mtafya, Ruth Bowness, Katarina Oravcova, Eoghan C W Farmer, Isobella Honeyborne, Dimitrios Evangelopoulos, Timothy D McHugh, Han Xiao, Celso Khosa, Andrea Rachow, Norbert Heinrich, Elizabeth Kampira, Gerry Davies, Nilesh Bhatt, Nyanda Elias Ntinginya, Sofia Viegas, Ilesh Jani, Mercy Kamdolozi, Aaron Mdolo, Margaret Khonga, Martin J Boeree, Patrick PJ Philips, Derek J Sloan, Michael Hoelscher, Gibson Sammy Kibiki, Stephen H Gillespie

## Abstract

**Objectives:** Better outcomes in tuberculosis require new diagnostic and treatment monitoring tools. In this paper we evaluated the utility of a marker of *M. tuberculosis* viable count, the Molecular Bacterial Load assay (MBLA) for diagnosis and treatment monitoring of tuberculosis in a high burden setting.

**Methods:** Patients with smear positive pulmonary tuberculosis from two sites in Tanzania and one each in Malawi and Mozambique. Sputum samples were taken weekly for the first 12 weeks of treatment and evaluated by MBLA and mycobacterial growth indicator tube method (MGIT).

**Results:** The results of high and low positive control samples confirmed inter site reproducibility. Over the 12 weeks of treatment there was a steady decline in the viable bacterial load as measured by the MBLA that corresponds to rise in time to a positive result (TTP) in the Mycobacterial Growth Indicator Tube. Both MBLA and MGIT provided similar time to test negativity. Importantly, as treatment progressed samples in MGIT were increasingly likely to be contaminated, which compromised the acquisition of results but did not affect MBLA samples.

**Conclusions:** MBLA produces a reproducible measure of *Mtb* viable count comparable to that of MGIT that is not compromised by contamination in a real-world setting. As a molecular test, the results can be available in as little as four hours and could allow health care professionals to identify rapidly patients who are failing therapy.

## Introduction

To help health care workers managing tuberculosis (TB) make better decisions we need to develop improved diagnostic methods and ways to monitor the response to treatment^1^. In addition to its clinical uses, a simple marker of treatment response would reduce costs in tuberculosis clinical trials and speed drug development ^2,3^.

Definitive diagnosis depends on the culture of *Mycobacterium tuberculosis* (Mtb), which is recognised as the current gold standard. For most TB control programmes there is, however, limited availability of culture due to the cost of high containment laboratories, staff training and reagent supply. With a culture-based approach results are available too late to inform timely decision making. They also produce variable results due to differences in specimen transport, media, decontamination method as well as the effect of bacterial or fungal overgrowth. In addition, sputum culture conversion is only weakly predictive of long-term patient outcome for regimens but not individuals^3^ limiting its application to individual and clinical trial management.

Enumeration of viable mycobacteria during therapy is probably the most accurate biomarker of treatment response currently available, but is technically difficult to standardise, expensive and results take many weeks to be reported^4^. Alternative more rapid methods include mycobacterial DNA-detection assays have established themselves as pre-treatment diagnostics, but prolonged DNA survival in the host after organisms have been killed precludes their use for treatment monitoring^5^. Messenger RNA (mRNA) targets, with a shorter half-life that DNA^7^ including *fbpB* antigen 85B, *hspX,* and *icl,* have been tested as treatment response biomarkers. Their decline during treatment mirrors traditional Mtb colony counting ^6–8^ but, as mRNA exists at low concentrations, mRNA based assays reach their limit of detection rapidly when patients are still TB culture positive ^7,8^. Consequently, such assays are valuable for following early responses, but monitoring over longer period is less effective.

The Molecular Bacterial Load Assay (MBLA), a real-time reverse transcriptase quantitative polymerase chain reaction (RT-qPCR) uses more abundant 16S-rRNA as a target and can accurately quantify Mtb viable bacillary load over many weeks of treatment^9^ and could replace solid culture^10^. We have now developed the assay further to provide enhanced stability for application in different settings including in high burden tropical countries^11^.

Here we report the first multi-centre evaluation of MBLA in comparison to standard methods in four high-burden settings in sub-Saharan Africa and we test whether it is reproducible, can assess disease severity and could provide a real time marker of a patient’s response to treatment.

## Methods

### Sites, patients and sample schedule

#### Tanzania

Patients were recruited from the PanACEA Multi-Arm Multistage (MAMS) TB trial sites Kibong’oto National Tuberculosis Hospital-Kilimanjaro Clinical Research Institute (KCRI) and National Institute of Medical Research at Mbeya Medical Research Centre (NIMR-MMRC)^12^. Tuberculosis was confirmed by sputum smear microscopy or GeneXpert. Multi-drug Resistant (MDR) TB was an exclusion and the experimental regimens are described in more detail in the MAMS study publication^12^. Early morning and spot sputum samples were collected at enrolment and weekly until 12 weeks. Culture was by Lowenstein Jensen (LJ) media and in the Mycobacterial Growth Indicator Tube (MGIT) liquid culture system and spot sputa were used for MBLA on site. If insufficient material was available the specimen was used for MGIT liquid culture.

#### Mozambique

Sputum smear and GeneXpert MTB/RIF positive patients, irrespective of HIV status were recruited from the Maputo Tuberculosis Trial Unit (MaTuTU). MDR-TB was not an exclusion criterion. Patients received WHO recommended treatment for susceptible or MDRTB as appropriate. Early morning and spot sputum samples were collected at enrolment and then one, two four, eight and 12 weeks.

#### Malawi

Sputum smear or GeneXpert MTB/RIF positive patients were recruited from the TB clinic, Queen Elizabeth Central Hospital, College of Medicine, Blantyre. All patients except two retreatment cases received standard treatment for susceptible disease. Early morning and spot sputa were collected at enrolment, then 2,4,6,8 and 12 weeks. LJ culture was only available for the first two weeks of treatment.

### Ethics

Consent for the collection and laboratory evaluation of samples from the MAMS-TB study was in accord with the clinical trial ethics approval, which was obtained from National Institute of Medical Research (NIMR) for the Tanzanian sites, NIMR/HQ/R.8c/242. Approval was also obtained from the Instituto Nacional de Saude (INS) Institutional Review Board and National ethics committee for the Mozambique site, 147/CNBS/14 and College of Medicine Research Ethics Committee (COMREC) University of Malawi for the Malawian site, P.08/13/1448.

### Molecular and Microbiological methods

Culture based measures: All sputum samples were tested by quantitative microscopy, culture on Lowenstein-Jensen medium and the Mycobacterial Growth Indicatory Tube as previously described ^12,13^.

#### Molecular Bacterial Load Assay (MBLA)

Sputum aliquots (1mL volume) were preserved for MBLA by diluting 1:4 in 50% guanidine thiocyanate (GTC) w/w, 0·1M Tris HCl pH 7·5 and 1% β-mercaptoethanol ν/ν then stored at - 80°C until testing. The internal control (100μΙ) was added to preserved sputum samples prior to RNA extraction and the mixture was centrifuged at 3000*g* for 30 minutes^11^. The tuberculosis molecular bacterial load assay was performed as described previously^11^.

The RT-qPCR was performed on site using a RotorGene 5plex platform (Qiagen). Primers and Taqman dual labelled probes targeting Mtb and the internal control were procured from Eurofin Genomics, Germany. A high (10^7^cfu/mL) and low (10^3^ cfu/mL) positive control (BCG NCTC 5692) and negative control of RNase free molecular grade water were included in each assay run.

#### Quality assurance and applicability of the MBLA assay in different laboratory environments

To verify the quality assurance, reproducibility and applicability of the different settings we analysed how participating laboratories processed the externally supplied standard BCG positive control panels. Two panels of material containing either a high concentration, 10^7^ CFU/ml and low concentration, 10^4^ CFU/ml (n=24) were supplied to all participating sites. Each laboratory independently conducted RNA extraction and qPCR and analysis.

### Analysis

Statistical analyses were performed using Graphpad Prism v.6, and the R statistical software environment (version 3.2.1). One-way analysis of variance (ANOVA) was applied to examine inter-site variance in MBLA performance. Sample “conversion” was defined as a change from ‘positive’ to ‘negative’ without subsequent reversion back to ‘positive’ before the end of follow-up. “Non-conversion” was defined as persistent or recurrently positive samples by the final visit on day 84. The day of conversion was defined as the midpoint between last positive and first definite negative result. A survival analysis was undertaken to investigate differences in time to conversion by assay, with data censored at 84 days (the time of last sample collection). Correlation of MBLA and MGIT was determined using Spearmans rank correlation (r). In all analyses p-values were considered significant at p<0.05

## Results

### Reproducibility

The results of the control samples tested in the four laboratories gave consistent measures of the bacterial load consistent with the spectrum of sputum samples tested: 7.1±0.3, 7.2±0.4, 7.3±0.6, 7.4±0.4 logio CFU/ml for the high concentration panel and 3.6±0.6, 3.6±0.7, 3.4±0.5, 3.6±0.4 for low concentration panel respectively. There was no significant variation in the measured bacterial load at the 4 sites, ANOVA p > 0.05 (Figure 1).

**Figure 1.**
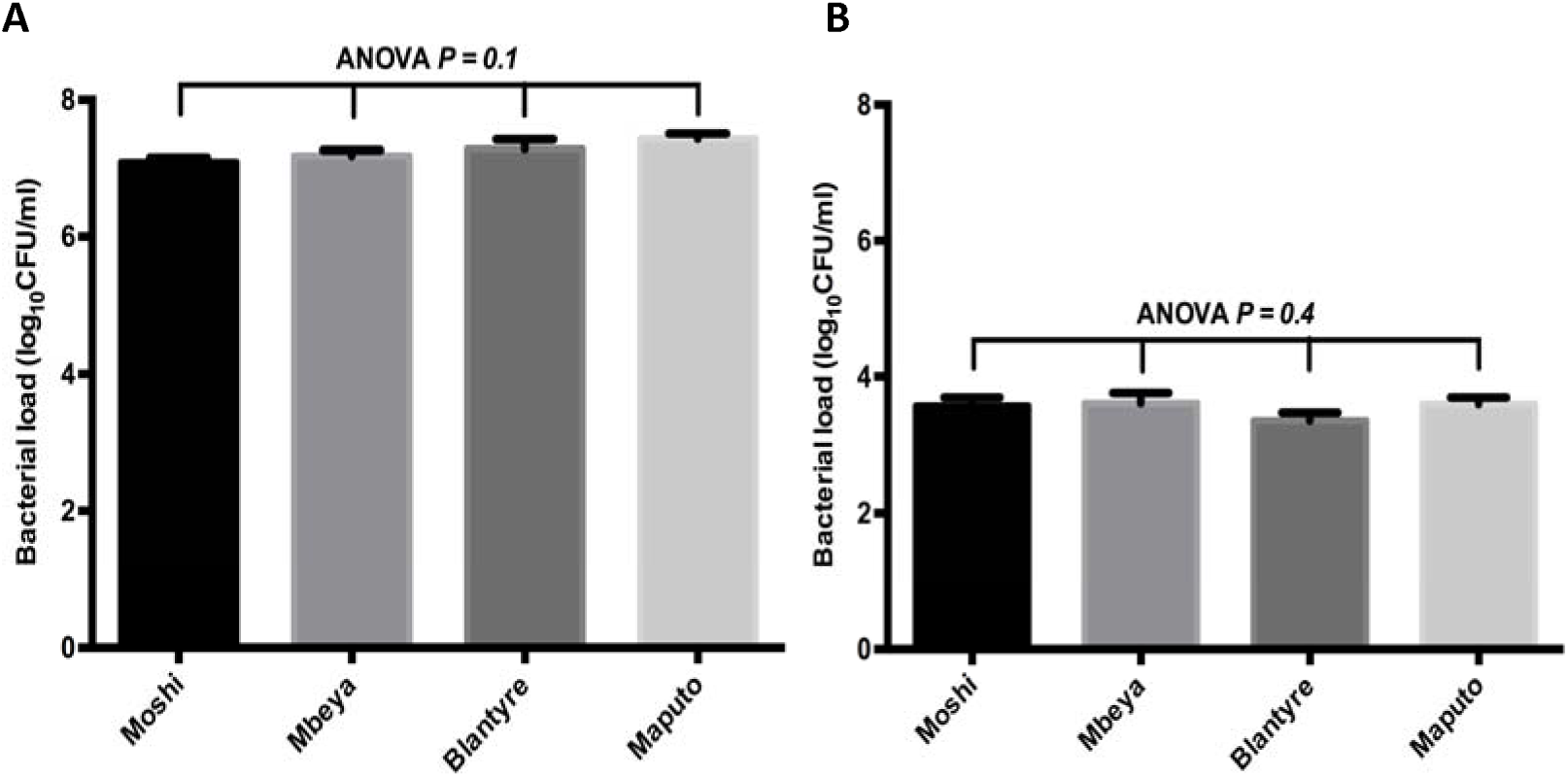
Comparison of MBLA control results in different laboratory settings. A) BCG(10^7^cfu/mL) high concentration panel (n = 24 per site) at the four sites, analysis of variance (ANOVA) *p = 0.1.* B) BCG (10^3^ cfu/mL) low concentration panel (n = 21 per site) at the four sites, ANOVA p =0.4 Error bars are standard error of the mean

### Patient recruitment

Of 213 anti-tuberculosis treatment naïve patients enrolled, 178 (83·6%) completed the follow-up and were included into the analysis: Tanzania 100 (55·9%), Mozambique 58 (32·8%), and Malawi 20 (11·3%). 128 (71·9%) were male and 47 (26·4%) were HIV positive with 33 years (IQR: 27-40 years) the median age. A further five patients (2.8%) were managed with an MDR regimen that included kanamycin, levofloxacin, ethionamide, cycloserine, and pyrazinamide. A total of 1768 serial samples were tested (Figure 2).

**Figure 2:**
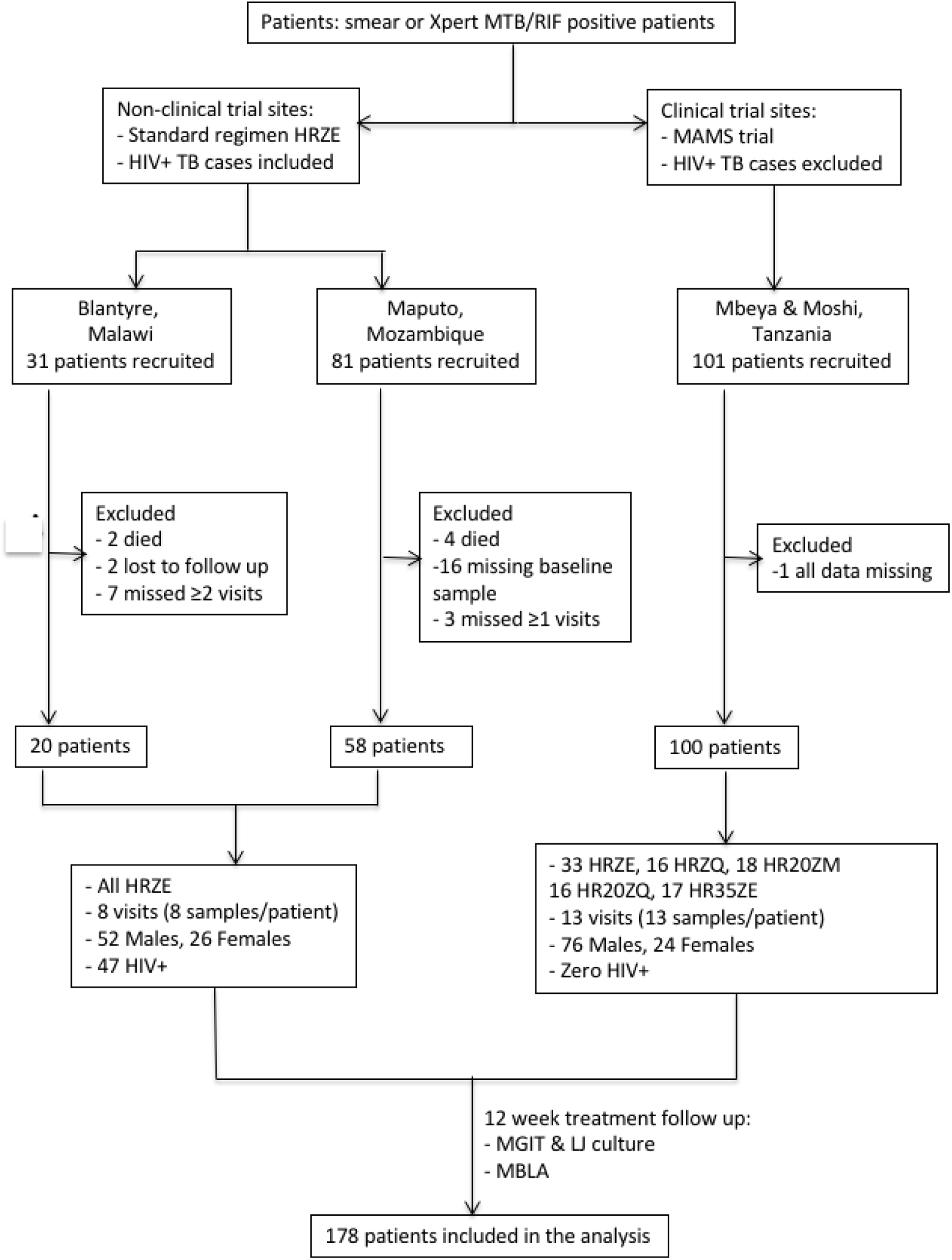
Enrollment and disposition of patients in the PANBIOME study

### Monitoring treatment response

We compared the assays on their ability to measure tuberculosis bacterial load. Taking the results from all of the patients with both measures at all time points we found that there was a strong correlation between the bacterial load as measured by MBLA and MGIT TTP, Spearmans r = −0.51, 95% CI (−0.56 to - 0.56), *p<0.0001* (Figure 3).

**Figure 3:**
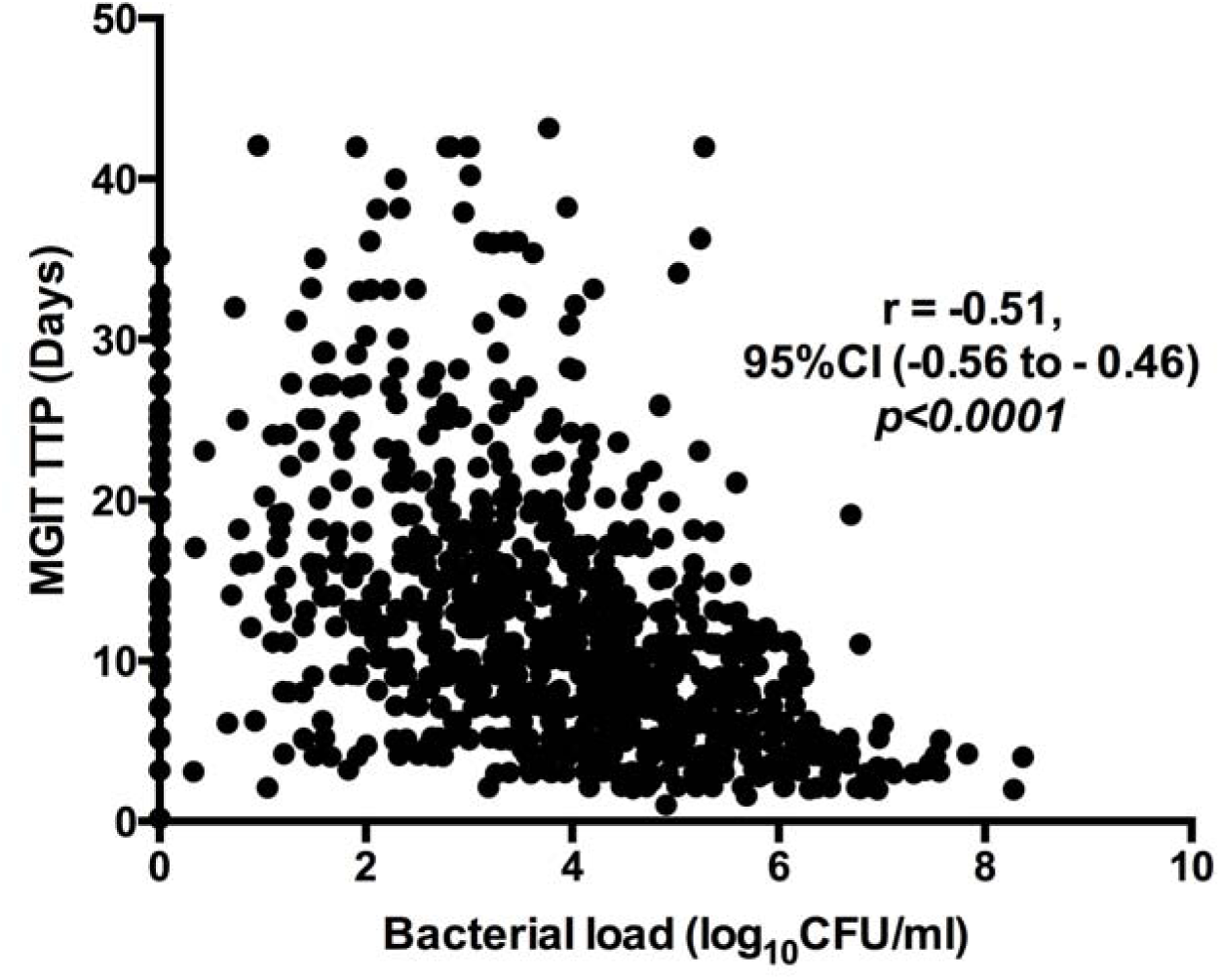
The Spearman’s rank correlation plot for correlation of bacterial load measured MBLA and TTP measured by MGIT. The higher the bacterial load the shorter the time to positivi

Defining treatment response as the fall in bacterial load measured by MBLA and or rising time to positivity (TTP) measured by MGIT we showed that, over the 12 weeks of treatment there was a steady decline in the bacterial load as measured by the MBLA that corresponded to a rise in time to a positive result (TTP) in the Mycobacterial Growth Indicator Tube as illustrated in Figure 4. Using the MBLA, the rate at which patient sputum cleared of TB was mean±SD, 1.09±l.llog_10_eCFU/ml/week and 0.84±0.6log_10_eCFU/ml/week by day 7 and 14 of treatment respectively. After day 14 the rate of sputum clearance plateaued at an average of 0.14±0.1log_10_eCFU/ml per week. The higher the bacterial load the steeper was the slope of the response curve in the first 2 weeks of treatment. Patients with above the average baseline bacterial load (5.53±1.3log_10_eCFU/ml) had significantly higher rate of clearance, 1.39±0.9log_10_eCFU/ml in the first 7 days and 0.99±0.5log_10_eCFU/ml in the 2^nd^ week of treatment compared to the low burden group clearance 1^st^ week 0.79±1.2log_10_eCFU/ml and 2^nd^ week 0.69±0.7log_10_eCFU/ml, Mann-Whitney test p<0.0001 and p=0.0002 respectively.

**Figure 4.**
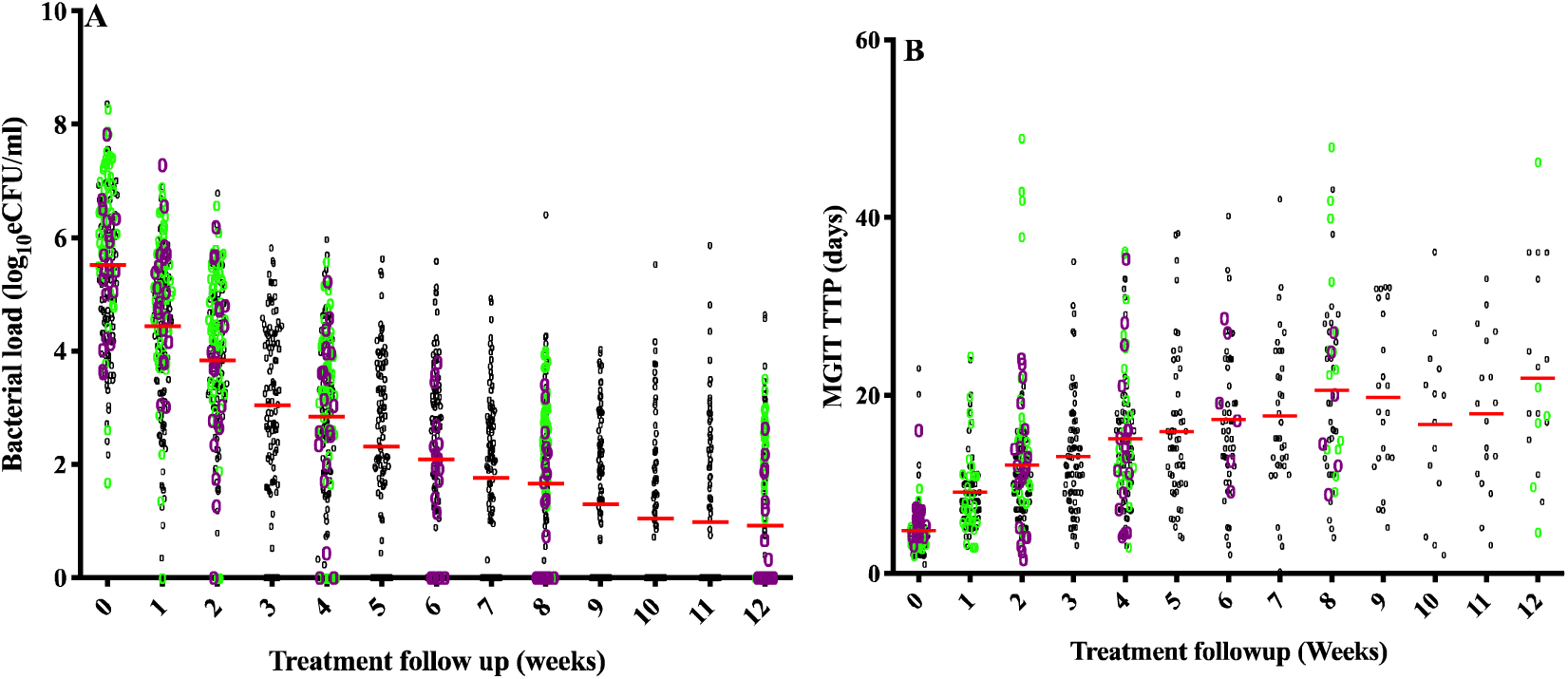
Bacteriological response to treatment measured by the Molecular Bacterial Load Assay (A) and by Mycobacterial Growth Indicatory Tube (B).Black = Tanzanian samples, Green+ Mozambique samples and Purple = Malawian samples Red lines represent the median values.

It is notable that the number of available points for MGIT reduces steadily as treatment progresses as samples are lost to contamination. In the first two weeks of treatment contamination rates are less than 10% but rise rapidly throughout the course of treatment until at week 12 more than half are lost to contamination.

We compared the qualitative performance of MBLA and MGIT culture during 12 weeks of treatment and these data are illustrated in Figure 5. At baseline, 171 (98%) patients generated quantitative MBLA eCFU/ml and MGIT TTP results. The number of dually positive samples decreased steadily as the number of positives declined and by week twelve only 16 (10·7%) were positive by both assays. A total of 232 (14·8%) samples were MBLA positive, MGIT contaminated including 32 (21·3%) from week 12. No patients were negative by MBLA and MGIT at baseline, but this figure rose to 20 (13·3%) by week 12. “MBLA negative, MGIT contaminated” was found in 195 (12·4%) specimens, including 61 (40·7%) at Week 12. In total, 427 (27·2%) samples with a definite “positive” or “negative” result on MBLA were lost to contamination by MGIT. Only 70 (4·5%) samples were MBLA negative and MGIT TTP positive (putative MBLA false negative). In comparison, 103 (6·57%) were MBLA eCFU/ml positive but MGIT negative (putative MGIT false negative). Solid (LJ) cultures, that were performed in Tanzania and Mozambique only, were less prone to contamination than MGIT at all time-points.

**Figure 5.**
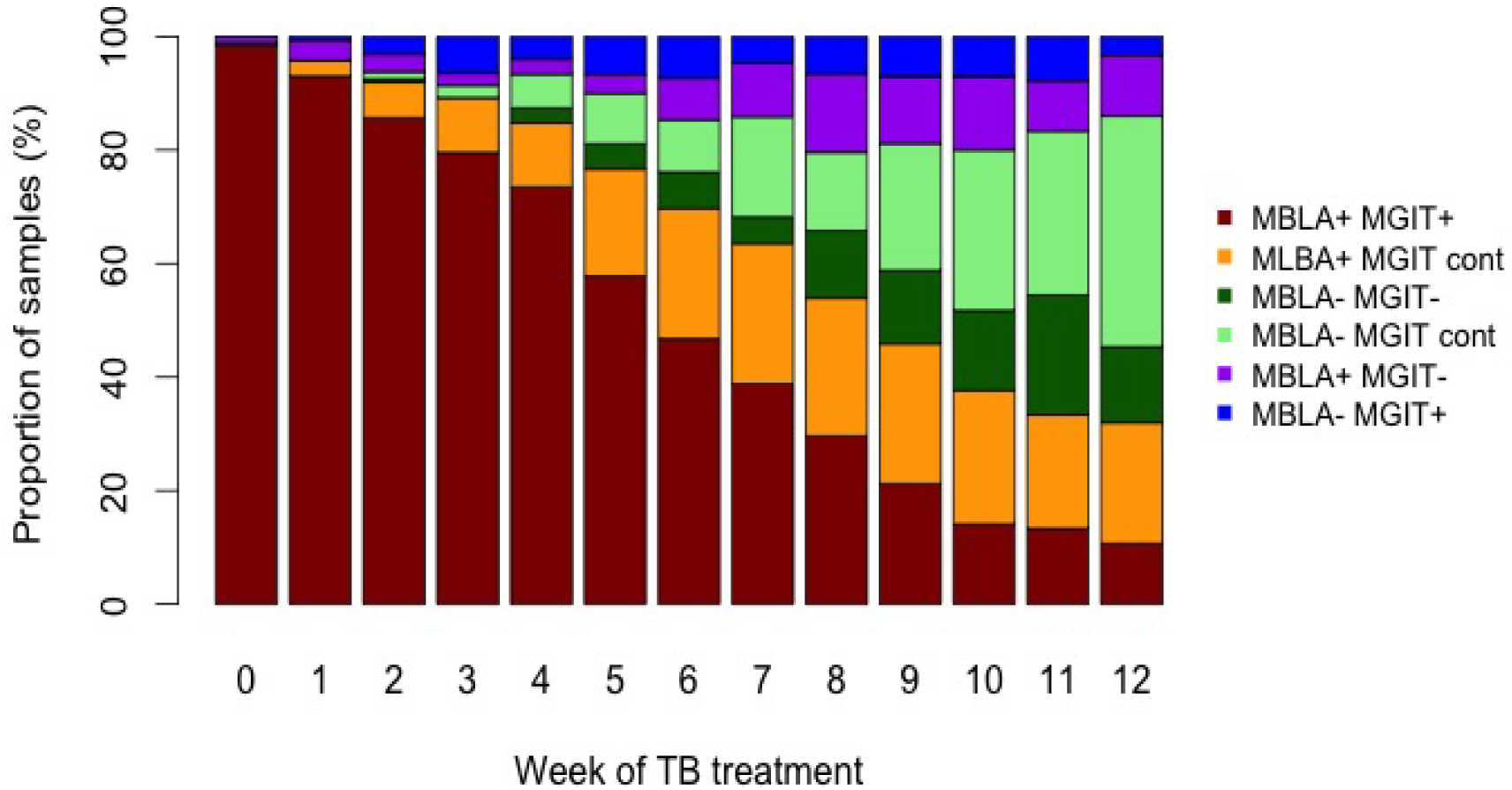
Comparison of the qualitative result (TB detected/not detected) by MGIT and MBLA and including the impact of contamination on the result from 171 patients with pulmonary tuberculosis.

We investigated how the MBLA, MGIT and LJ, assessed conversion to negativity in this population. In those with susceptible organisms, median time to conversion was similar on MGIT and MBLA but was shorter on LJ. It was notable that almost half of the MGIT and LJ results were undeterminable due to contamination (Table 1).

**Table 1:** Comparison of proportion of negative, contaminated and missing results at different time points for MBLA, MGIT and LJ and the time to negativity by these three measures. UD = Undetermined data point (i.e contaminated, neither positive nor negative). Conversion to negativity results are mean ±SD of the data set.

| Form of TB | Status | MBLA | Liquid culture | Solid culture (LJ) |
| --- | --- | --- | --- | --- |

**Table 1: Comparison of proportion of negative, contaminated and missing results at different time points for MBLA, MGIT and LJ and the time to negativity by these three measures.**
|  |  |  | (MGIT) |  |
| --- | --- | --- | --- | --- |
| **Susceptible (N=164)** | Positive Week 0 (%) | 100 | 96 | 57 |
|  | Positive Week 8 (%) | 68 | 23 | 14 |
|  | Positive Week 12 (%) | 41 | 10 | 4 |
|  | UD Week 0 (%) | 0 | 3 | 28 |
|  | UD Week 8 (%) | 0 | 65 | 54 |
|  | UD Week 12 (%) | 0 | 70 | 69 |
|  | Missing Week 0 (%) | 0 | 0 | 0 |
|  | Missing Week 8 (%) | 0 | 5 | 0 |
|  | Missing Week 12 (%) | 0 | 20 | 0 |
|  | Time to conversion (days) | 66±21 | 65±22 | 34±21 |
| **MDR (n=4)** | Positive Week 0 (%) | 100 | 100 | 75 |
|  | Positive Week 8 (%) | 75 | 0 | 0 |
|  | Positive Week 12 (%) | 100 | 25 | 25 |
|  | UD Week 0 (%) | 0 | 0 | 0 |
|  | UD Week 8 (%) | 0 | 75 | 50 |
|  | UD Week 12 (%) | 0 | 25 | 50 |
|  | Missing Week 0 (%) | 0 | 0 | 0 |
|  | Missing Week 8 (%) | 0 | 25 | 0 |
|  | Missing Week 12 (%) | 0 | 0 | 0 |
|  | Time to conversion (days) | 84±00 | 42±28 | 39±33 |
| **Monoresistant (N=7)** | Positive Week 0 (%) | 100 | 100 | 43 |
|  | Positive Week 8 (%) | 29 | 0 | 0 |
|  | Positive Week 12 (%) | 14 | 0 | 14 |
|  | UD Week 0 (%) | 0 | 0 | 14 |
|  | UD Week 8 (%) | 0 | 43 | 14 |
|  | UD Week 12 (%) | 0 | 43 | 0 |
|  | Missing Week 0 (%) | 0 | 0 | 0 |
|  | Missing Week 8 (%) | 0 | 0 | 14 |
|  | Missing Week 12 (%) | 0 | 14 | 14 |
|  | Time to conversion (days) | 47±28 | 55±21 | 29±19 |
| **Polydrug resistant (N=3)** | Positive Week 0 (%) | 100 | 100 | 33 |
|  | Positive Week 8 (%) | 33 | 33 | 0 |
|  | Positive Week 12 (%) | 67 | 67 | 0 |
|  | UD Week 0 (%) | 0 | 0 | 67 |
|  | UD Week 8 (%) | 0 | 33 | 67 |
|  | UD Week 12 (%) | 0 | 33 | 67 |
|  | Missing Week 0 (%) | 0 | 0 | 0 |
|  | Missing Week 8 (%) | 0 | 33 | 0 |
|  | Missing Week 12 (%) | 0 | 0 | 0 |
|  | Time to conversion (days) | 63±25 | 84±0 | 40±15 |

## Discussion

The most important questions facing health care workers managing patients with tuberculosis is to confirm the diagnosis and to determine whether patient are responding to the prescribed therapy. Failure to respond may be due to multiple factors including poor adherence poor absorption of drugs, or a drug-resistant infecting organism. Current diagnostic methodologies do not provide information to address this crucial clinical question in a meaningful time-frame. In this paper, we report the results of a large-scale multi-centre field trial to further assess the applicability of the tuberculosis MBLA in high burden settings^9^.^10^

All of the sites established the assay successfully and when their performance was compared using control samples similar results were obtained. This confirms that the test results are reproducible in a high-burden setting. We have shown previously that TB MBLA is species specific confirming the diagnosis ^11 14 9^ and this provides the healthcare worker with confirmation of the tuberculosis diagnosis.

We show that, for our population as a whole there is an inverse correlation between MBLA and TTP. When plotted as a patient journey, the decline in TB MBLA is an inverse image of the MGIT TTP results over the first three months of treatment^9^. This suggests that they are monitoring broadly similar, clinically relevant measures in the viable count. In the first few weeks following the initiation of treatment, it is important to understand whether the patient is responding. A failure to respond may be due to the presence of a resistant strain, patient non-adherence, poor drug absorption or counterfeit medicine. Current diagnostic tests do not provide the healthcare worker with data to detect poor response. Radiological appearances do not resolve in a timely or reproducible way^15^. MGIT is a semi-quantitative measure of viable count as the patient responds to therapy the time to a positive result increases, and the time taken to report a useful result increases proportionately whether positive or negative.

We have shown that MBLA is at least as sensitive as GeneXpert although MBLA does not provide simultaneous resistance determination. MBLA targets a species of nucleic acid, rRNA that is more stable than mRNA and present in higher concentration. Detecting the presence of rRNA indicates that the organism is viable^9^.^10^. Thus, MBLA delivers data similar to culture-based methods: a viable count with patients having between one hundred and one hundred million organisms per mL at baseline. These numbers are very similar to other studies allowing for the impact of different culture media^4,13,16^, confirming previous studies showing the utility of using MBLA to predict the pretreatment bacterial burden.

The quantitative read out provided by MBLA is an advantage as it provides a direct measure of bacterial load. This provides a significant advantage over Lowenstein Jensen medium or on the MGIT where the results are only be semiquantitative and patient progress cannot be judged readily. From the perspective of the treating healthcare worker, the MBLA provides a readily understandable result a number of viable bacteria compared to time to positivity.

An important advantage of using the MBLA is that the results are not compromised by bacterial overgrowth. Sputum samples contain many bacteria other than the target mycobacteria and, since these grow more rapidly, their growth must be suppressed. This is achieved by decontaminating the sample with bactericidal chemicals such as sodium hydroxide followed by the incorporation of malachite green in Lowenstein Jensen and a cocktail of antibiotics in MGIT. Both of these processes reduce the bacterial load and introduce an opportunity for variation in the laboratory performance and result. Contaminated samples are more likely as the patient’s treatment progresses and they have greater difficulty producing a sample reducing its overall quality whilst at the same the number of mycobacteria falls that may be adversely affected by decontamination^17^. In our data we show the impact of this effect with significant loss of samples due to bacterial contamination that becomes more significant as treatment progresses. As the MBLA result depends on species specific primers it is unaffected by the presence of other bacteria which means that as treatment progresses MBLA almost always is able to produce a quantitative value for the *Mtb* viable count whereas and increasing number of MGIT samples are lost.

Despite the positive data reported in this paper, there are a number of issues that still need to be addressed. Our study included patients who were infected with susceptible organisms and all responded well to therapy. Further evaluation is needed in many more patients with resistant disease, a setting where MBLA may have a lot to offer. It is envisaged that to assess treatment response two samples would be collected, on commencing therapy and after a period treatment. Poor responders would be identified by the failure of the bacterial load to fall. This begs two unanswered questions: when is the optimal time for the second sample and what is a bacterial load decline that implies a successful or unsuccessful outcome? These questions will be addressed in clinical studies that are ongoing. Although we have shown here that MBLA can generate consistent results in different centres in high burden settings the assay still requires molecular expertise and a well-equipped laboratory in a research setting. There is a need to simplify the assay to so that it is easy to perform with minimal hands on time and available at affordable cost.

In summary, we have shown that the TB MBLA is a reproducible test of *M. tuberculosis* viable bacterial load with potential for diagnosis that can be implemented successfully in a high burden setting. It is rapid to perform and is not affected by bacterial contamination. It can deliver data on the number of viable bacteria in as little as four hours and it could be used to assess initial severity of disease and monitor the response to treatment. Further work is required to simplify the assay to make it accessible and develop the correct sampling schedule to achieve the assay’s full potential

